# A generative network model of neurodevelopment

**DOI:** 10.1101/2020.08.13.249391

**Authors:** Danyal Akarca, Petra E Vértes, Edward T Bullmore, the CALM team, Duncan E Astle

## Abstract

The emergence of large-scale brain networks, and their continual refinement, represent crucial developmental processes that can drive individual differences in cognition and which are associated with multiple neurodevelopmental conditions. But how does this organization arise, and what mechanisms govern the diversity of these developmental processes? There are many existing descriptive theories, but to date none are computationally formalized. We provide a mathematical framework that specifies the growth of a brain network over developmental time. Within this framework macroscopic brain organization, complete with spatial embedding of its organization, is an emergent property of a generative wiring equation that optimizes its connectivity by renegotiating its biological costs and topological values continuously over development. The rules that govern these iterative wiring properties are controlled by a set of tightly framed parameters, with subtle differences in these parameters steering network growth towards different neurodiverse outcomes. Regional expression of genes associated with the developmental simulations converge on biological processes and cellular components predominantly involved in synaptic signaling, neuronal projection, catabolic intracellular processes and protein transport. Together, this provides a unifying computational framework for conceptualizing the mechanisms and diversity of childhood brain development, capable of integrating different levels of analysis – from genes to cognition.

---

Macroscopic brain organization is present early in life, even prenatally^1,2^, and continues to develop through adolescence and into early adulthood^3^. The emergence and continual refinement of large-scale brain networks, connecting neuronal populations across anatomical distance, allows for increasing functional integration and specialisation^4,5^. This process is thought crucial for the growth of complex cognitive processes such as language^6^ and executive function^7–12^. However, there are individual differences in the trajectories of these networks, and these mirror differences in cognitive development. Indeed, differences in macroscopic networks have been implicated across multiple neurodevelopmental conditions^13^, including ADHD^14^, autism^15,16^, and language disorders^17^.

But what mechanisms govern the growth of macroscopic brain networks? And how do these mechanisms give rise to individual differences in developmental outcomes? There are numerous descriptive theories of typical and atypical neurodevelopment^18–22^ that speculate about how different levels of analysis (e.g. genes, brain structure and function) interact to guide development. However, to date there are no *generative* models of neurodevelopment. That is, no existing theories are sufficiently specified that they can simulate individual-level brain networks. In the absence of computational models of development, it is difficult to establish mechanistic links between different observations (e.g. genes, biological pathways, system wide organization). This theory gap represents a major limitation for developmental systems neuroscience. The purpose of this study is to address precisely this gap, by modelling the generative wiring properties of the developing brain, guided by a simple principle: the brain’s structural organization is shaped by an economic trade-off between minimising wiring costs and adaptively enhancing valuable topological features^23^. We hypothesize that the emergence of whole-brain organization reflects the continual trade-off of these factors over developmental time and that tiny differences in the parameters governing the trade-off can produce the neurodiverse outcomes we observe. Somewhat counterintuitively, tight parameter constraints likely enable macroscopic neurodiversity, because large changes in these parameters would produce networks with configurational states that are not observed in reality. Instead, narrow boundaries reflect parameter conditions within which networks can be different, but still maintain adequate structural properties to be functional.

We used generative network modelling^24,25^ in an intentionally heterogeneous sample of children (N=270), and: (i) tested which topological features should be valued in the growth trade-off to produce highly accurate individual child connectomes; (ii) tested how small changes in these growth parameters alter the organizational properties of the resulting networks; (iii) established relationships between these different growth parameters and cognitive development; (iv) identified genes with expression profiles that were spatially colocated with those topological features; and (v) established the biological pathways that are enriched in these gene lists. Together, this provides a computational framework that mathematically specifies the growth of a network over developmental time, captures individual differences in brain organization and cognition, and incorporates the genetic and biological pathways that likely constrain this development.

## RESULTS

### The generative network model

The generative network model (GNM) can be expressed as a simple wiring equation^19,20^ (**Fig. 1a**), which formalizes a trade-off between costs and adaptive value when brain regions form connections probabilistically over time:

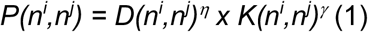

**Figure 1.**
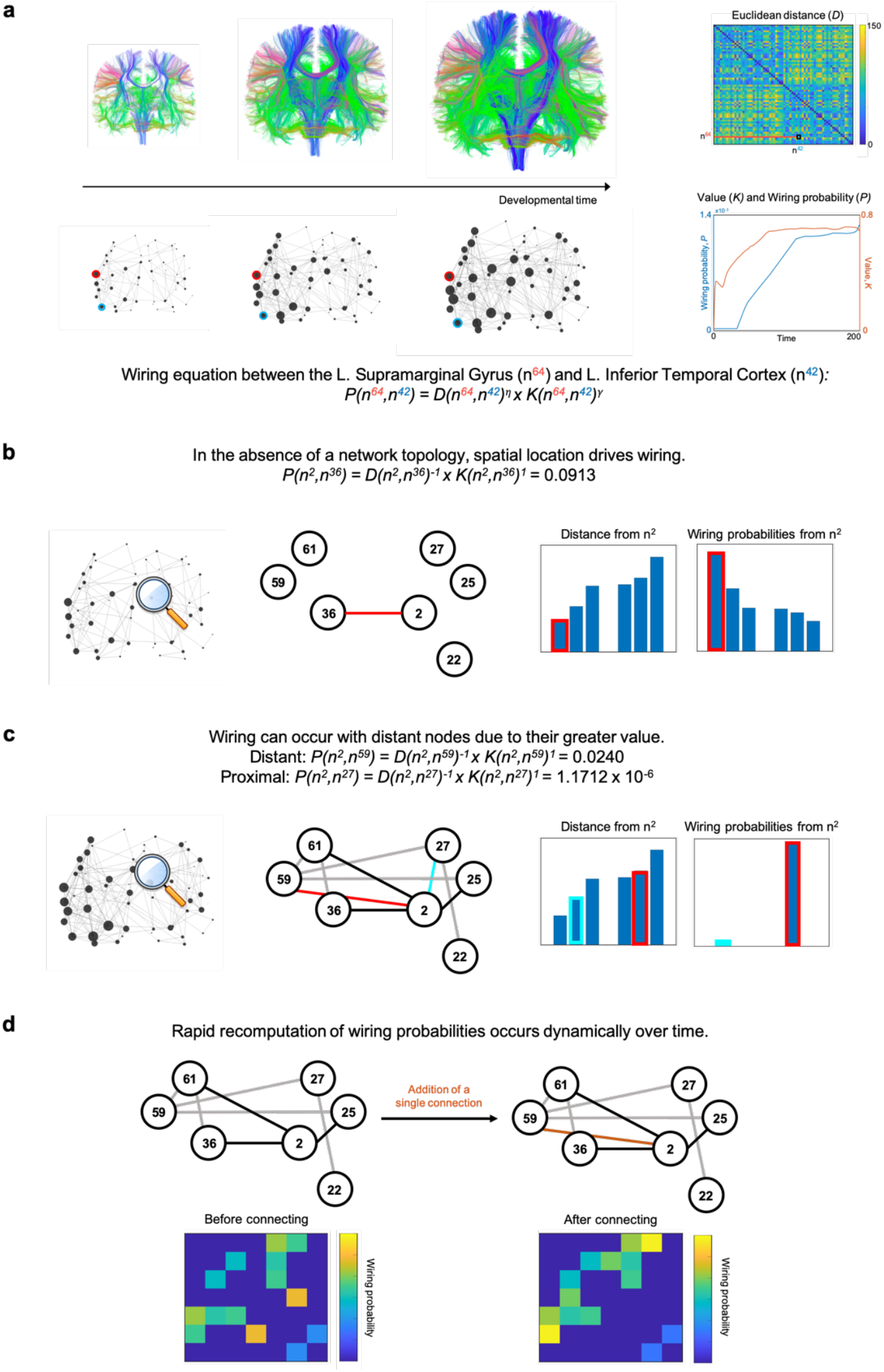
Updating wiring probabilities within the generative network model iteratively, based on dynamically changing graphical structures. (**a**) The brain’s structural connectivity is modelled as a generative network which grows over time according to parametrized connection costs, *D*(*n^i^,n^j^*)^*η*^, and values, *K*(*n^i^, n^j^*)^*γ*^. In this illustration, we use subject one’s optimal model. (**b**) Early in network development, the absence of a topology leads to proximal nodes being much more likely to form connections. The displayed distances and probabilities are from the right caudal anterior cingulate (n^2^), which corresponds to row *D*(*n*^2^,:) and *P*(*n*^2^,:). We display it’s six nearest cortical regions. (**c**) Later, the relative values (*K*) between nodes influence connection probabilities, such that nodes which are more distant (e.g. left rostral anterior cingulate, n^59^ in red) may be preferred to nodes which are closer (e.g. right superior frontal cortex, n^27^ in cyan). (**d**) As costs and values are decoupled, the wiring probability can be rapidly recomputed when dynamic changes in graphical structure occur over developmental time.

Where *D*(*n^i^, n^j^*) represents the Euclidean distance between nodes *n^i^* and *n^j^* (i.e. ‘costs’), and *K*(*n^i^, n^j^*) reflects the value (i.e. ‘attractiveness’) in forming a connection. *P*(*n^i^, n^j^*) represents the wiring probability resulting from the product of the parameterized costs and value.

*D* is parameterized by the scalar *η*, which influences how node-to-node distances influence their probability of connecting. For example, when *η* is negative, wiring probabilities decline when distances increase, and this reflects the costliness of forming connections with nodes that are distant. This is traded-off against *K*, which represents some relationship between nodes. *K* can be thought of as a topological value (or ‘rule’) driving the intention for node *n*^i^ to connect with node *n^j^*. *K* is parameterized by a distinct scalar *γ. K* can take a range of different forms and can, in principle, be selected from any non-geometric growth rule used to model social and economic networks^26–29^. One simple example is the ‘matching’ rule^24^: nodes form connections with other nodes on the basis of their normalized overlap in neighborhood – i.e. whether nodes are connected to similar nodes to themselves (also termed *homophily*).

Imagine the following scenario: nodes in a network are growing according to the matching rule, preferentially attaching to nodes which are both similarly connected and spatially proximal. In the wiring equation, this would be represented as *η* being negative (e.g. *η* = −1), *K* represented as normalized neighborhoods between nodes (i.e. matching) and its parameter *γ* being positive (e.g. *γ* = 1). In short, a node being far away makes it less likely that a new connection will be formed, but it having a similar a neighborhood increases the likelihood. Suppose that the right caudal anterior cingulate (Node 2, *n*^2^) is going to wire to one of its six nearest neighbors. Initially, due to an absent network topology, spatial proximity has a great influence in the formation of new connections – it will be wired to its nearest neighbor (**Fig. 1b**). However, gradually over time, the network’s developing structural topology means that *K* (i.e. the relationships between nodes) may now have a greater influence on wiring probabilities. Indeed, the right caudal anterior cingulate may later wire with a node that, although it is further away than other available nodes, has a greater value (i.e. matching) than the others (**Fig. 1c**). As the wiring equation separately parameterize costs and value, the presence of a single connection can heavily influence the topology of the network and thus the wiring probabilities. This is because new connections can lead to entirely new overlapping neighbors, which may include distant nodes. As a result, wiring probabilities can change considerably from moment to moment, despite costs remaining fixed (**Fig. 1d**).

The GNM simulates this process across the whole brain, until the overall number of connections matches those in the real brain network. Subsequently, to test whether the model has produced a network that closely approximates real brain networks, an energy function, *E*, must be defined which measures the dissimilarity between simulated and observed networks^24,25^:

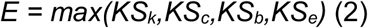

Where *KS* is the Kolmogorov-Smirnov statistic comparing degree *k*, clustering coefficient *c*, betweenness centrality *b* and edge length *e* distributions of simulated and observed networks.

Minimising *E* finds GNM parameters *η* and *γ* which generate networks that most closely approximate the observed network.

### Small variations in GNM parameter combinations produce accurate and spatially embedded networks

We computed the subject-wise optimal GNM (i.e. network with lowest energy) over a range of 10,000 evenly-spaced parameter combinations (*η* ± 7, *γ*± 7) using thirteen different generative rules (for rule formulae, see **Supplementary Table 1**) across our large sample of children (*N* = 270, 178 males, 92 females, mean age = 9yrs 10m, SD age = 2yrs 2m). In each case, we computed energy landscapes to contextualize how they perform based on their class (e.g. degree) and computation (e.g. product) (**Fig. 2a–2d**). Mirroring previous findings in adult samples^24,25,29^, we found that models driven by geometry and topology outperform the pure geometric spatial model and homophily-based models achieve the lowest energy for our pediatric sample (**Fig. 2e**). In other words, when one combines the distance penalty with the ‘matching rule’ we described in our example (as shown in **Fig. 1**), it produces the most plausible brain networks.

**Figure 2.**
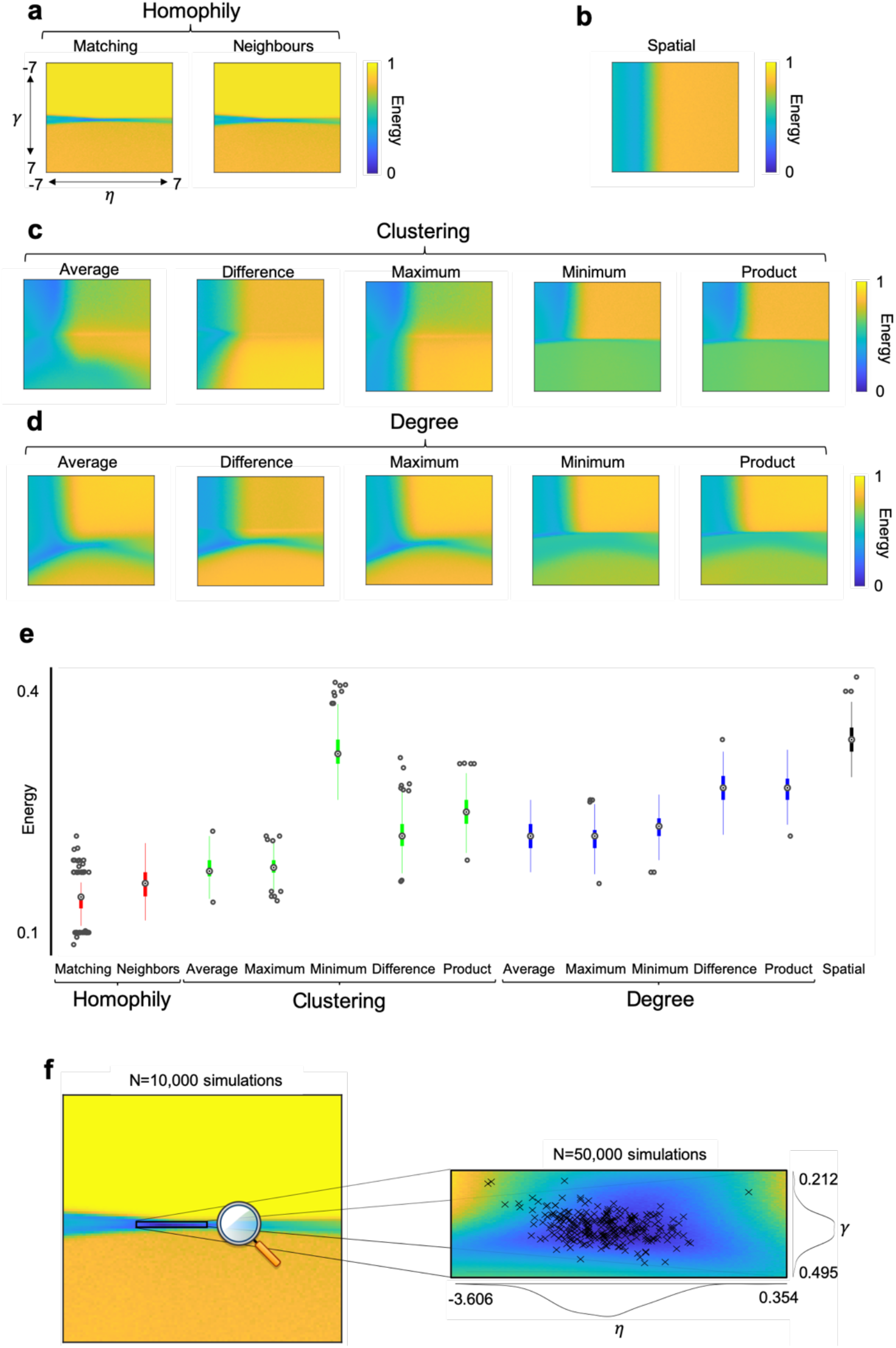
Sample-averaged energy landscape visualization and generative rule comparisons. (**a**) Homophily based methods. Matching and neighbors algorithms calculate a measure of shared neighborhood between nodes. **(b**) The spatial method. This ignores *γ* entirely, judging networks only on the basis of their spatial relationship. (**c**) Clustering based methods. These calculate a summary measure between two nodes in terms of their clustering coefficients. (**d**) Degree based methods. These calculate a summary measure between two nodes in terms of their degree. (**e**) Energy statistics from the best performing simulation across generative rules, showing that matching achieves lowest energy networks. A tabulated form of this figure is provided in **Supplementary Table 1**. (**f**) A further 50,000 simulations were undertaken in the refined matching window, as these defined boundary conditions for which low-energetic networks were consistently achieved. Each cross represents a subject’s individually specific wiring parameters that achieved their lowest energy simulated network. Energy coloring is taken from group averages.

It is notable that across the matching energy landscape, the most plausible networks exist within an extremely narrow parameter window. That is, as a proportion of the parameter space, the matching rule (and the other homophily-based model ‘neighbors’) contain the least number of low-energy networks relative to other rules. But as **Figure 2e** shows, these networks are the closest to real networks. Thus, varying homophily-based parameters produces the most realistic networks, yet has the lowest variability in the space.

While small, variability within this narrow matching window determines inter-individual differences in brain network growth. This is because small changes in parameters (i.e. the magnitude and direction in which costs and values influence wiring probabilities) can lead to networks which are diverse yet include basic structural properties common to all participants. To derive more precise estimations of optimal generative parameter combinations, we subsequently generated a new set of 50,000 evenly-spaced simulated networks over this narrow low-energy matching window (−3.606 < *η* < 0.354, 0.212 < *γ*< 0.495). Focusing on this energy crevasse allows us to detect individual differences in optimal parameter combinations with much greater specificity.

Within this refined window, we calculated optimal wiring parameters for each subject in our sample. In **Figure 2f**, we show the spatial distribution of top performing parameter combinations and **Supplementary Table 2** documents their summary statistics. Intuitively, lower-energy networks were found than before due to a much higher resolution of parameter selection. In **Supplementary Figure 1a-d** we detail how KS statistics vary across the same space.

Due to the stochastic nature of GNMs, the energy of optimal parameter combinations varies with an average standard deviation (SD) of 0.045 across the sample (1000 independent runs). Therefore, for the rest of this study we quote our parameter analyses averaged across a variable number of wiring parameters which achieved networks with the lowest energy in the space: N = 1 (equating to 0.002% of the space) N = 10 (0.02%), N = 100 (0.2%) and N = 500 (1.0%). We find that optimal *η* and *γ* parameters are significantly negatively correlated with each other, such that subjects with large *γ* parameters tend to have larger negative *η*(Best N=1 network: R = −0.284, P = 2.07 x 10^-6^; N = 10 networks: R = −0.403, P = 6.08 x 10^-12^; R = −0.460, P = 1.58 x 10^-15^; N = 100 networks: R = −0.460, P = 1.58 x 10^-15^, N = 500 networks: R = −0.497, P = 3.21 x 10^-18^) (**Supplementary Fig. 1e**). Optimally simulated networks, using this simple wiring equation, are so similar to the actual networks that a support vector machine is unable to distinguish them using the parameters from the energy equation (2) (mean accuracy = 50.45%, SD = 2.85%).

Simulated networks are optimized using the energy equation (2), and the best performing networks capture the statistical properties of the observed networks (**Fig. 3a-d**)^24,25^. But will the best networks also capture crucial network properties not included in the energy equation, like the spatial embedding of the statistical properties within the network? We next examined if the spatial patterning of topological and network characteristics spontaneously arises from the constraints of the wiring equation.

**Figure 3.**
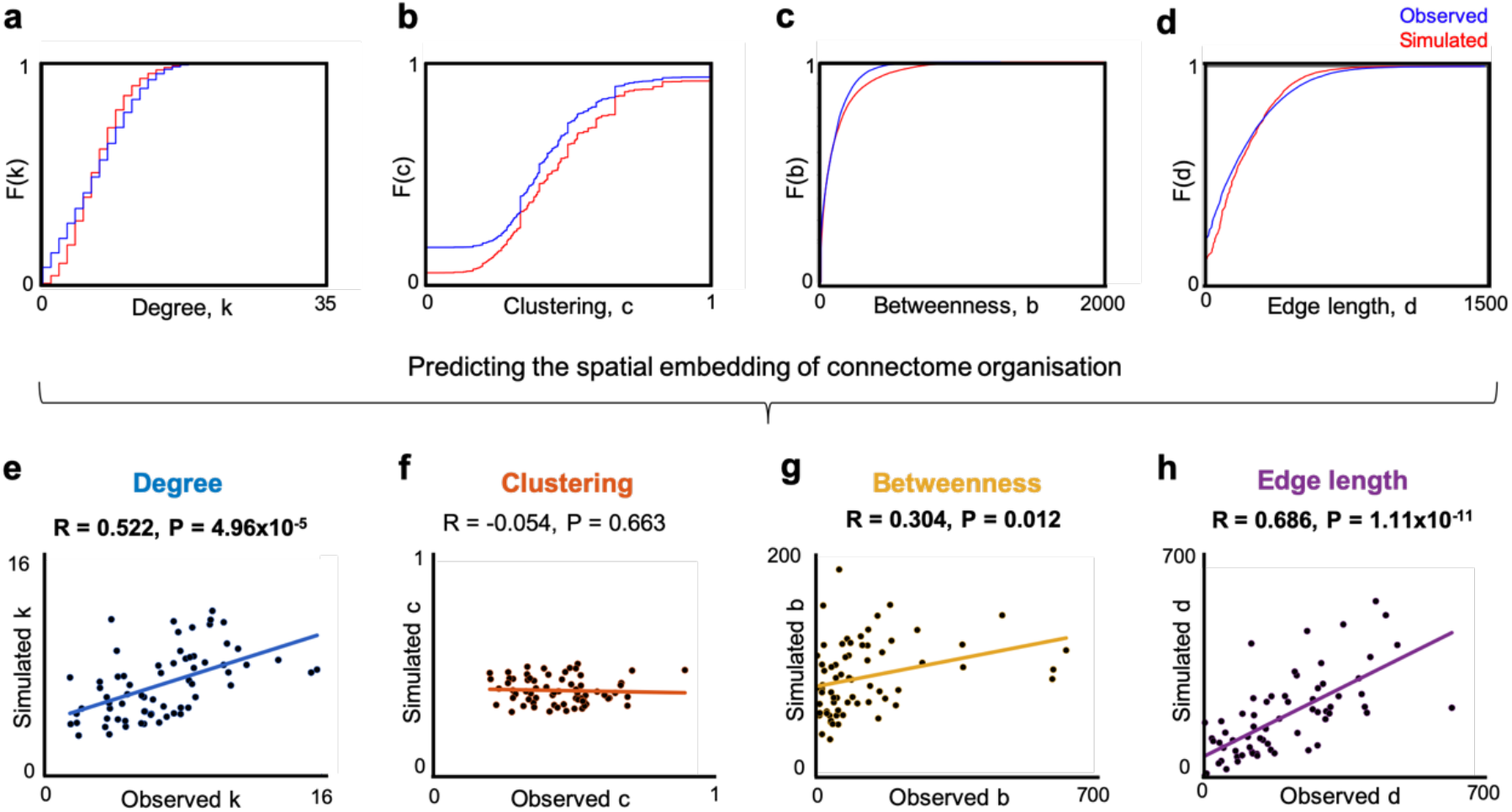
Cumulative density functions of topological and network features included in the energy equation and predictions of their spatial embedding. Each point in the scatter is the across-subject average nodal measure from the observed and optimal simulated networks in terms of (a) Degree (b) Clustering coefficient (c) Betweenness centrality (d) Edge length. (e-h) Linear correlations between observed and optimally simulated networks across the sample for each measure. Supplementary Figure 2 shows a parallel analysis for measures outside of the equation. Boldened values are significant correlations at P<0.05.

Across the sample, optimally performing generative models (i.e. those using the ‘matching’ rule) produce networks which significantly correlate with observed networks in terms of their degree (R = 0.522, P = 4.96×10^-5^), edge length (R = 0.686, P = 1.11×10^-11^) and betweenness centrality (R = 0.304, P = 0.012) but not clustering coefficient (R = −0.054, P = 0.663) (**Fig. 3e-h**). That is, the spatial embedding of these network properties spontaneously emerges, to mirror those of the observed networks, despite this not being specified in the growth process. We extended this analysis to new measures outside of the energy equation (**Supplementary Fig. 2**). While local efficiency and assortativity cannot be significantly predicted across the sample (R = 0.211, P = 0.084 and R = −0.096, P = 0.116 respectively), optimally performing simulated and observed networks correlate positively in terms of their global number of rich clubs (R = 0.316, P = 1.11×10^-7^), maximized modularity (R = 0.349, P = 3.84×10^-9^) and transitivity scalar (R = 0.411, P = 2.11×10^-12^). In short, this simple growth rule also generates many of the crucial properties of real brain networks, despite them not being specified in the growth process.

### Individual differences in parameters mirror connectome organization and morphology

So far, wiring equations need to incorporate a ‘matching rule’ to best approximate real brain networks. Tiny changes in the wiring parameters are needed to best approximate individual connectomes across our diverse sample. But do these differences in wiring parameters influence structural brain organization and do they reflect any observed features of a subject’s brain? We next quantified how a subject’s *η* and *γ* individually relate to global measures of their observed connectome. Furthermore, for all 270 subjects, cortical morphology data were also available. In **Figure 4** we show that a subject’s *η* and *γ* are associated with a range of global network and morphological measures, both inside and outside of the energy equation. This is an important step in validating that these parameters generalize to distinct measures (e.g. morphological observations) not used to train the generative models.

**Figure 4.**
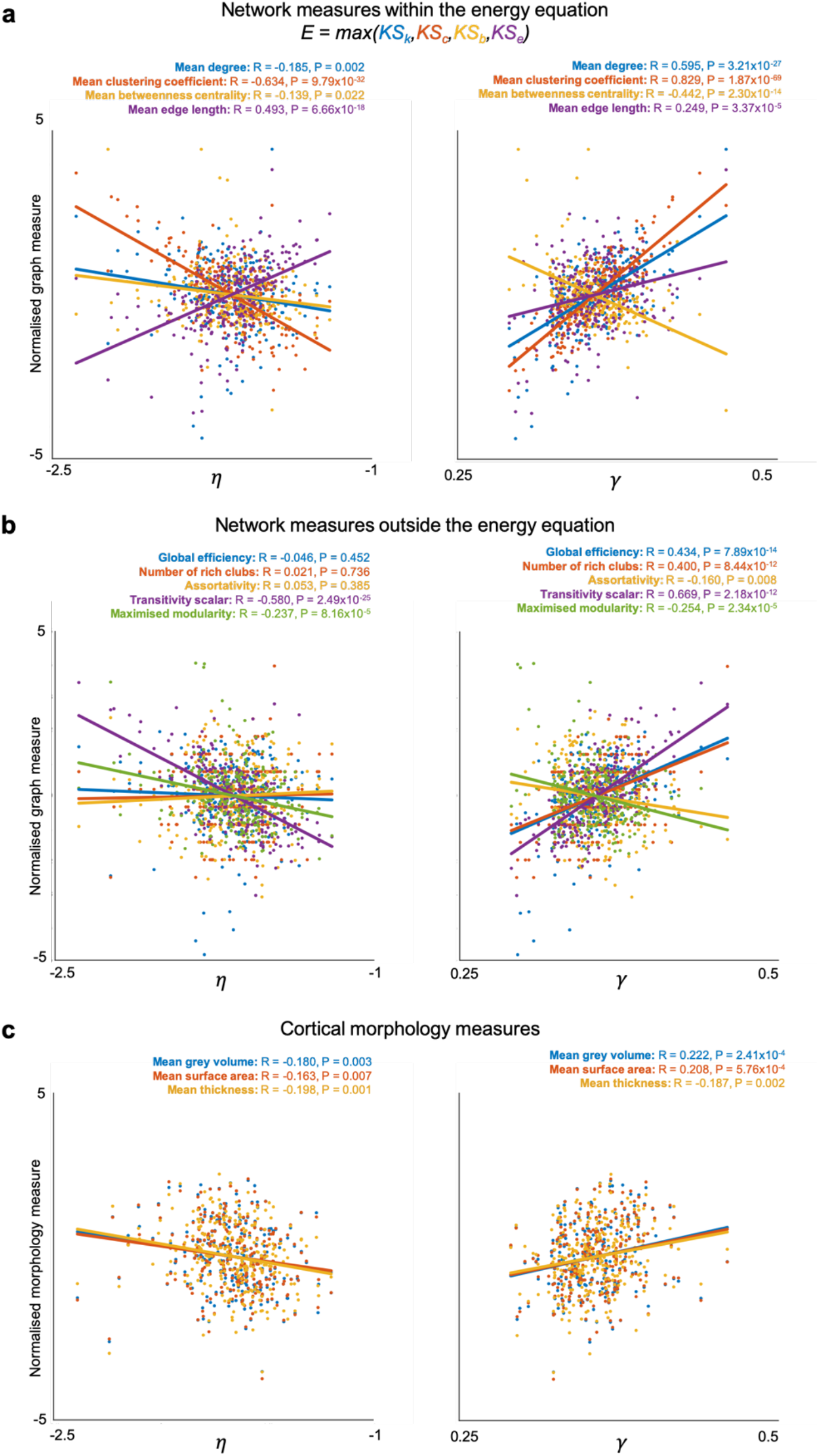
Variations in a subject’s wiring parameters *η* and *γ* relate to global structural properties of their brain networks and their cortical morphology. These parameters were calculated by averaging across the top N = 500 simulated networks, but findings for variable N networks are provided in **Supplementary Table 3**. (**a**) *η* and *γ*significantly relates to all measures in the energy equation. (**b**) *η* relates to transitivity and modularity, but not efficiency, number of rich clubs or assortativity all of which were not minimized in the energy equation. *γ* relates to all examined measures not included in the energy equation. (**c**) *η* is negatively correlated with morphological findings, while *γ* is positively correlated with morphological findings.

Across subjects, *η* correlates negatively with mean degree (R = −0.185, P = 0.002), global clustering (R = −0.634, P = 9.79×10^-32^), mean betweenness centrality (R = −0.139, P = 0.022) and positively with mean edge length (R = 0.493, P = 6.66×10^-18^). *γ* correlates positively with mean degree (R = 0.595, P = 3.21×10^-27^), global clustering (R = 0.829, P = 1.87×10^-69^), mean edge length (R = 0.249, P = 3.47×10^-5^) and negatively with mean betweenness centrality (R = −0.442, P = 2.30×10^-14^) (**Fig 4a**). Despite not being explicitly minimized, *η* correlates negatively with transitivity (R = −0.580, P = 2.49×10^-25^) and modularity (R = −0.237, P = 8.16×10^-5^). It does not correlate with global efficiency (R = −0.46, P = 0.452), the number of rich clubs (R = 0.021, P = 0.736) or assortativity (R = 0.053, P = 0.385). By contrast, *γ* correlates positively with global efficiency (R = 0.434, P = 7.89×10^-14^), number of rich clubs (R = 0.400, P = 8.44×10^-12^), transitivity (R = 0.669, P = 2.179×10^-12^) and negatively with assortativity (R = −0.160, P = 0.0083) and modularity (R = −0.254, P = 2.34×10^-5^) (**Fig 4b**). To summarize briefly: at a global level, the tiny amounts of variability in these two parameters across the children significantly predict a wide range of properties of their observed connectomes.

Finally, we tested for this correspondence in a totally different type of neuroimaging data – cortical morphology. *η* correlates negatively with mean cortical grey matter (R = −0.180, P = 0.003), surface area (R = −0.163, P = 0.007) and thickness (R=-0.198, P = 0.001). By contrast, *γ*is positively correlated with mean cortical grey matter (R = 0.222, P = 2.41×10^-4^), surface area (R = 0.208, P = 5.76×10^-4^) and thickness (R=0.187, P = 0.002) (**Fig. 4c**). These global findings are stable, regardless of how many high performing parameters are averaged across, and they are all outlined in **Supplementary Table 3**.

### Variability in neurodevelopmental trajectories arises through value-updating over time and relates to cognitive outcomes

While small generative parameter differences result in differential network properties, we have yet to show how this variability occurs over developmental time. That is, how do differences in parameter combinations across subjects manifest themselves when the brain network is developing? To address this, we examined how between subject-variability in optimal GNMs emerge at the level of cortical nodes and their connections. This is possible by simply decomposing the optimal simulation into its constituent parametrized costs (*D^η^*), values (*K*) and wiring probabilities (*P*) at each time point, for each subject (**Fig. 5a-b**). This allows us to quantify growth trajectories and thus establish which aspects of network emergence vary most in the sample.

**Figure 5.**
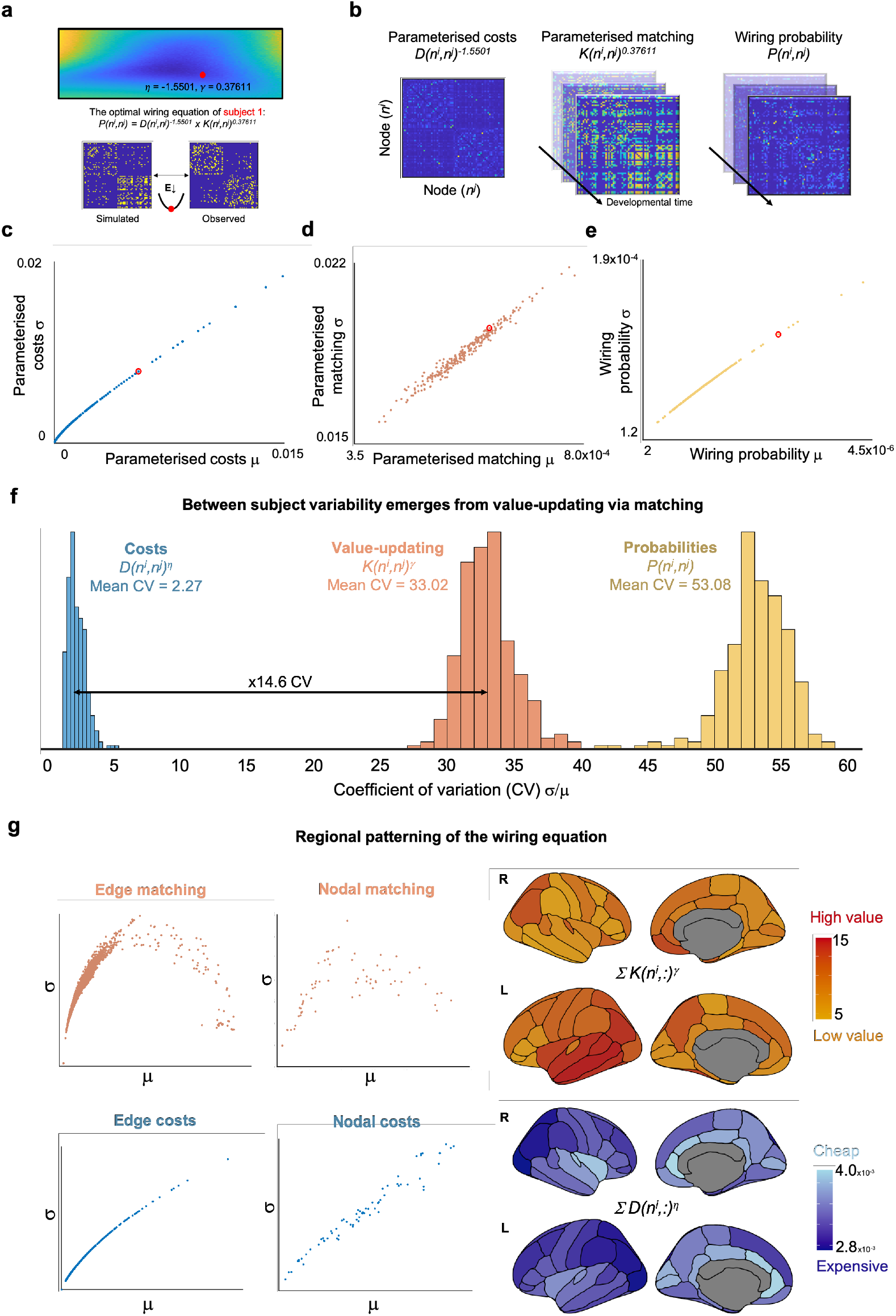
Wiring equation (1) decomposition and the subsequent variability across subjects in our heterogeneous sample. (**a**) For each subject, a simulated network is produced by minimizing the energy between the observed and simulated network. Here we present visualizations for subject one (red). (**b**) Costs (*D*) are static, while values (*K*) dynamically update according to the matching rule, which enables the computation of wiring probability (*P*). (**c)** The mean and standard deviation for each subject of their edge-wise parameterized costs, (**d**) parameterized matching and (**e**) wiring probabilities. (**f**) Histograms of each subject’s coefficient of variation (CV) showing that subjects are more variable in theirs valueupdating compared to costs, which leads to large wiring probability variation. (**g**) Regional patterning of sample-averaged nodal parameterized costs and values, showing highly valuable patterning in the left temporal lobe and cheap regions generally occupying medial aspects of the cortex. Variability declines as value increases, but increases for costs.

For each subject, we computed the coefficient of variation (CV, σ/μ) of their parameterized costs, matching values and wiring probabilities to compare subject-specific variability, as it emerges throughout the simulated growth of connectomes. This allows for a comparison in variability between components of the wiring equation. While subjects exhibit some variability in how parameterized costs influence wiring probabilities (mean CV 2.27), this is dwarfed by their parameterized matching over time (mean CV 33.02). This is because the matching value is dynamic, changing at each iteration (as in **Fig. 1d**) unlike relative Euclidean distance between nodes, which is static. The result is that significant inter-individual variability arises in the probability of connections forming (mean CV 53.08), leading to the emergence of divergent brain organization (**Fig. 5c-f**). Furthermore, the regional patterning of costs and values is not random (**Fig. 5g**). Nodes and edges with high matching values decline in their variability, suggesting a consistency across subjects in highly ‘attractive’ nodal structures and their connections. Across the sample, cheaper regions occupy the medial aspects of the cortex while highly valuable regions generally reside in the left temporal cortex.

We have shown that the complex architecture of a child’s connectome can emerge simply by estimating the changing wiring properties over its development, generalizing to multiple different properties of their observed connectomes and cortical morphology data. But do these growth parameters relate to their cognition? If wiring parameters are indeed accurate decompositions of an individual’s structural network, we should find that they predict cognitive outcomes equivocally to observed features of the connectome. For all 270 subjects we had data from a battery of cognitive tasks, including measures of executive function, phonological awareness, working memory, fluid reasoning and vocabulary.

We tested the relationship between a subject’s cognition and (i) their optimal wiring parameters and (ii) global measures of their structural connectome (using measures included in the energy equation). This was done using partial least squares (PLS), a multivariate statistical technique which extracts optimally covarying patterns from two data domains^30^. We undertook two separate PLS analyses, which correlated (i) optimal wiring parameter combinations or (ii) global connectome measures across our sample, with cognitive performance in the nine tasks, respectively (**Fig. 6a**). For both analyses, PLS1 was significant in the amount of explained covariance (76.2% explained, p_corr_ = 0.009 and 62.6%, p_corr_ = 0.049 respectively). PLS1 score predictions, and their cognitive loadings, are extremely similar between wiring parameters and connectome features (R = 0.191, P = 1.63 x 10^-3^ and R = 0.210, P = 5.29 x 10^-4^) (**Fig. 6b-c**).

**Figure 6.**
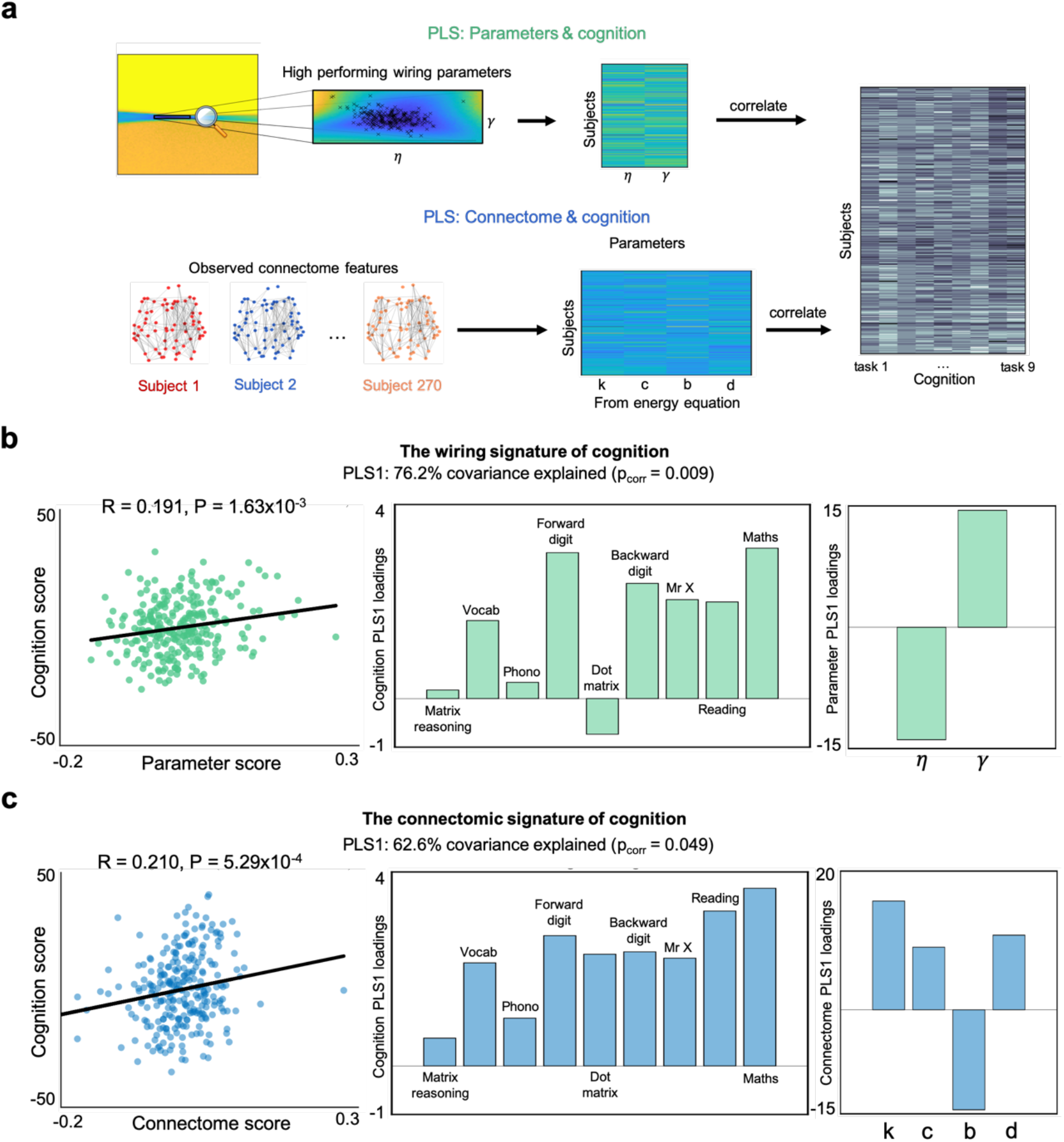
Partial least squares (PLS) regression was used to identify covarying patterns of wiring parameters and global connectome features with cognitive performance across nine cognitive tasks. (**a**) A visual representation of the two PLS analyses undertaken. PLS1 was statistically significant (p_corr_ = 0.009 and p_corr_ = 0.049 respectively) for both analyses, accounting for 76.2% and 62.6% of the covariance between predictors and cognition respectively. (**b**) There is a significant positive correlation between parameter scores and PLS-derived cognitive scores. Each parameter loads with similar magnitude onto PLS1 (note that *η* is a negative value) (**c**) There is an analogous significant positive correlation between connectome scores and PLS-derived cognition scores.

### Genomic patterning of network growth

Underlying these macroscopic changes in brain organization across time are a series of complex molecular mechanisms. These are partly governed by genetically coded processes that vary across individuals. We next tested whether these processes may steer the brain network towards a particular growth trajectory within our GNMs.

Nodal cost and nodal ‘matching’ value patterning alongside regional gene expression profiles of 10,027 genes using human adult brain microarray data^31,32^ were integrated into two PLS analyses for each subject. For all analyses, gene expression scores at each node were used as the predictor. For each subject’s first analysis, their parameterized nodal costs (calculated as *D^η^* as visualized for subject one in **Fig. 4b; left**) was used as the response variable. For each subject’s second analysis, their mean parameterized values (calculated as *K^γ^* averaged over time, as visualized for subject one in **Fig. 4b; middle**) was used as the response variable. Each analysis defined PLS components independently which were linear combinations of the weighted gene expression scores at each node (predictor variables) that were most strongly correlated with the subject’s nodal costs and nodal values of their simulated growth trajectory. To limit the variability across regions in terms of the samples available, only left hemispheric gene data were analyzed.^27^

Across our sample, the first PLS component (PLS1) explained on average 65.0% (SD 1.3%) and 56.9% (SD 9.2%) of the covariance between genetic expression and nodal-costs, and nodal-values respectively. The average nodal-costs PLS1 score significantly correlates with average nodal-costs (R = 0.794, P = 2.07×10^-8^). Similarly, the average nodal-values PLS1 score significantly correlates with average nodal-values (R = 0.718, P = 1.71×10^-6^) (**Fig. 7a-b**).

**Figure 7.**
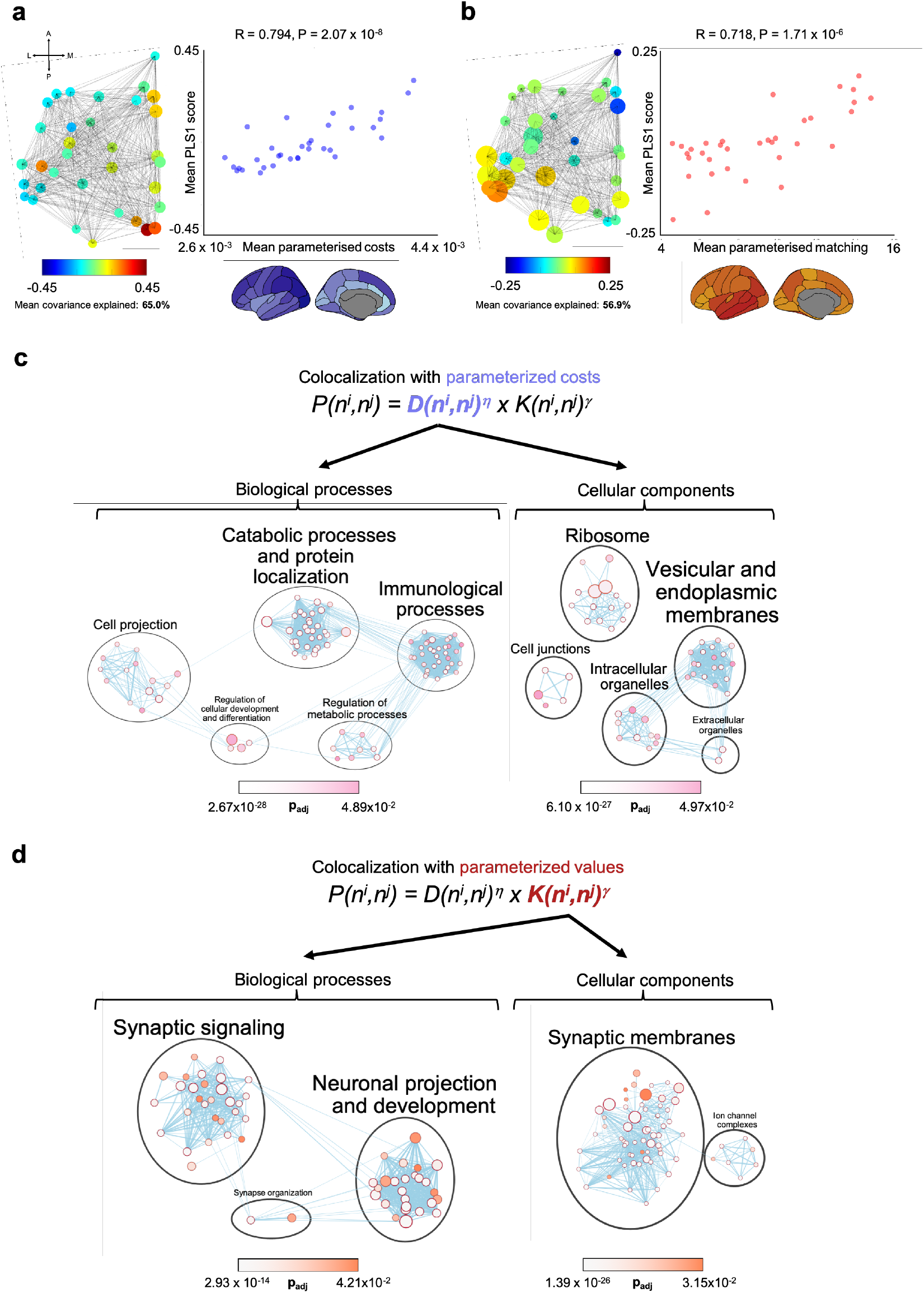
PLS regression identifies over expressed genes which explain variance in parameterized costs and values across subjects. Both PLS1 components across subjects are enriched for functionally specific biological processes and cellular components. Node size represents the number of genes in the set. The edges relate to gene overlap. (**a**) Sample averaged parameterized costs significantly correlates with sample averaged PLS1 nodal gene scores, explaining on average 65.0% covariance (**b**) Sample averaged parameterized values significantly correlates with sample averaged PLS1 nodal gene scores explaining on average 56.9% covariance (**c**) Nodal-costs PLS1 is enriched for genes predominantly associated with protein localization, catabolic processes and ribosomal/membrane cellular components (**d**) Nodal-values PLS1 is enriched for genes predominantly associated with synaptic signaling, neuronal projection and synaptic membranes.

To then characterize the genetic profiles associated with each PLS analysis, we permuted the response variable 1000 times to form a null distribution for the loading of each gene, across each subject’s PLS1. This provides an estimate of how strong the loading would be by chance, and thus which genes exceed p_corr_<0.05. Across subjects, PLS1 provided an average of 581.5 significant genes (SD 101.4) for nodal costs and 437.6 significant genes (SD 167.4) for nodal values (**Supplementary Fig. 3a**).

Genes do not act in isolation, but instead converge to govern biological pathways across spatial scales. To move from individual genes to biological processes (BPs) and cellular components (CCs), we performed a pathway enrichment analysis^33^. Pathway enrichment analysis summarizes large gene sets as a smaller list of more easily interpretable pathways that can be visualized to identify main biological themes. Genes were ordered according to their frequency in being significantly associated with connectome growth across subjects for that component. For example, for nodal-values PLS1, top of the list was the gene associated with connectome growth in the most subjects (*CHI3L1*; significant for 49.4% of our sample), the next was the second most frequent gene (*PRKAB2*; 36.4% of our sample) and so on. Our list stopped when genes were significant for less than 10% of the sample. This left the nodal-costs PLS1 with a list of 1427 genes and the nodal-values PLS1 with a list of 1584 genes ordered in terms of importance, which were submitted to pathway enrichment analysis (**Supplementary Fig. 3b**).

The genes identified within the subject-wise PLS are not random, but instead converge on particular biological processes (BP) and cellular components (CC). The nodal costs PLS1 was most prominently enriched for genes associated with biological processes including catabolic processes and protein localization (32 BPs; all p_adj_ < 9.58 x 10^-3^), cell projection (14 BPs; all p_adj_ < 4.39 x 10^-2^), immunological processes (34 BPs; all p_adj_ < 4.82 x 10^-2^), regulation of metabolic processes (8 BPs; all p_adj_ < 4.75 x 10^-2^) and regulation of cell development and differentiation (4 BPs; all p_adj_ < 3.87 x 10-2). In terms of cellular components, nodal costs PLS1 was enriched for genes associated with the ribosome (14 CCs; all p_adj_ < 2.15×10^-2^), vesicular and endoplasmic membranes (19 CCs; all p_adj_ < 4.90 x 10^-2^) and intracellular organelles (8CCs; all p_adj_ < 4.97 x 10^-2^) (**Fig. 7c**).

The nodal values PLS1 was most prominently enriched for genes associated with biological processes including synaptic signaling (29 BPs; all p_adj_ < 3.96 x 10^-2^), neuronal projection and development (26 BPs; all p_adj_ < 4.21 x 10^-2^) and synapse organization (2 BPs; all p_adj_ < 2.92 x 10^-2^). In terms of cellular components, nodal values PLS1 was enriched for genes associated with synaptic membranes (60 CCs; all p_adj_ < 3.15 x 10^-2^) and ion channel complexes (7 CCs; all p_adj_ < 1.18 x 10^-2^) (**Fig. 7d**).

In **Supplementary Table 4** we provide links so that readers can run our precise gene ontology queries within a browser and in **Supplementary Figure 3c-d** we show a visualization of these enriched gene sets.

## DISCUSSION

The principles of the development of macroscopic human brain organization can be modelled as a generative network, that optimizes its connectivity by renegotiating its costs and value^23,24^ continuously over time. Despite the simplicity of this equation, it results in the dynamic updating of wiring probabilities over development, with multiple network properties, like spatial embedding, being an emergent property of this dynamic updating. This resonates with theoretical perspectives that implicate dynamic interactions between brain systems over development in progressive, integrative, specialisation^34^. We have formalized this process in the context of neurodevelopmental diversity; offering a new perspective on the emergence of macroscopic organization, its possible biological underpinnings, and the association with functional outcomes like cognitive performance. This reflects a theoretical step-change in the study of human brain development, in being sufficiently well-specified to generate macroscopic brain networks. In turn, this formalization allows for the unpacking of the computational and/or biological constraints that shape the trajectories of networks. Indeed, we anticipate that GNMs may be a powerful tool to model the growth and diversity of real and biologically-feasible artificial networks across many scales.

Small changes in growth parameters of the GNM lead to divergent macroscopic brain networks, with systematically different network properties. Within the model, the key factor that drives individual differences in growth trajectory is the dynamic nature of updating preferences. Specifically, as nodes form new connections this dynamically changes their neighborhoods, and in turn this quickly changes which nodes become ‘attractive’ for subsequent connections. Importantly, individual differences in this process correspond significantly to independent structural data of the same individuals.

Why do the homophily-based generative rules approximate whole-brain networks so well? We propose that the superordinate goal of any developing brain network is to achieve the optimal computational capacity required of it, given finite biological resources. In this light, we suggest that matching produces the lowest-energetic networks precisely because it provides the closest *heuristic* estimate (compared to those tested here and in other works^24,25,28^) of the genuine dynamic reappraisals that occur over developmental time. This is because by virtue of preferentially wiring with nodes with shared neighborhoods, modular architectures emerge, and this reflects the brain’s overarching structure. The modular architecture of the brain has been well studied, and has numerous properties enabling effective flexible computations likely important for functional integration^35^. Finally, by virtue of only requiring knowledge of neighborhood overlap, homophily-based methods may incur less informational costs^36^ relative to other methods which require global information, and therefore may be more biologically plausible.

Our current GNMs operate at a whole-brain level – i.e. a global set of rules governing growth. But this could likely be made more biologically realistic by integrating regional differences that capture the potential long-term benefits of forming a connection. For instance, methodological improvements might be achieved by integrating reinforcement learning models with GNMs. This would model regional brain structures as agents aiming to optimize their connectivity dynamically over time to maximize their long-term reward. Many superficial strategies could be used to achieve this, and the appearance of the network’s “growth rules” would likely change considerably at different time points to achieve the superordinate goal. For example, while in earlier stages rewards may best be attained via the rapid generation of the modular network, once a sufficient architecture is produced it may be better attained via its remodeling to reduce maintenance costs^23^. Such a model may also incorporate wider factors, such as environmental influences, which may influence reward predictions and thus affect the steering of the network’s trajectory. This conceptual understanding of network growth complements the wider literature surrounding “computationally rational”^37^ navigational strategies (such as the successor representation^38^ or intuitive planning^39^) that can be taken by agents in a dynamic environment under constraints. This area of study has mainly focused on understanding how agents can represent and optimize navigation through cognitive maps to attain some reward (e.g. such as accumulating rewards in a maze). However, the application of these approaches may prove useful to understand how networks themselves optimize their connectivity at different stages of development.

Regional variation in nodal costs and values closely mirrored the expression profiles for different sets of genes, which in turn govern different biological processes and cellular components. Since the advent of genome-wide association studies (GWAS), a huge number of genes have been implicated in developmental disorders, including schizophrenia^40^ and autism^41^, but also general cognitive functioning^42^. It has been challenging to interpret the consequences of these individual implicated genes. The enrichment analysis that accompanied our GNM takes a very different approach. As far as we are aware, this is the first study aiming to bridge models of whole brain organizational emergence and genetics in this way (for work using generative models, see refs^24,25,29,43,44^ and work that integrates Allen Brain Atlas gene data with functional and structural brain imaging, see refs^45–48^). Nodal costs covaried with genes enriched for highly costly metabolic processes, including catabolic processes, protein transport and cellular components centered around the ribosome and endoplasmic membranes. On the other hand, nodal values covaried with those enriched for trans-synaptic signaling, neuronal projection and the synaptic membrane.

The *omnigenic* model^49^ suggests that complex traits are driven by genes that do not have direct effects on the trait per se, but instead propagate through regulatory networks on much smaller numbers of core genes, with more direct effects. This model explains the vast number of GWAS hits for complex traits, as “peripheral” genes necessarily outnumber “core” genes and thus the sum of their small effects exceeds the contribution of core genes. We suggest the omnigenic model may apply to some aspects of gene-development relationships. That is, the many genes that contribute to each PLS1 may not directly contribute to developmental processes themselves, but in the regulation of activity and growth within brain areas that are particularly important for neurodevelopment. Crucially there is variability in enriched genes across subjects (**Supplementary Fig. 3a**).

This work presents a challenge to the developmental field which has a long history of categorizing neurodevelopment into discrete groupings based on observed cognitive and/or behavioral traits. Instead, we suggest that neurodevelopment may be better thought of as a trajectory in which divergent outcomes arise via slight trajectory changes that fall out of the continual negotiation of brain connectivity optimization. While likely that generative preferences are initialized via an individual’s genetic preprograming, small changes in wiring preferences over time – possibly via complex interactions of their time course, endocrinological exposure, learning and environment – have profound effects on the emergence of the developmental trajectory. What results is a continual interaction between network growth preferences and the dynamically developing brain, leading to neurodiverse outcomes.

This computational framework has a number of limitations that provide scope for future improvements. Our generative models are limited to the binary connections which are assumed to be anatomical^24,25^. This is inevitably a gross simplification of the complex weighted structure of the connectome. Devising ways in which network connections can change in a more graded fashion is a necessary next step to modelling more complete developmental processes. In the future we will need to capture both the strengthening and weakening of connections that has been shown to occur in human brain development^50,51^. Secondly, we currently use one rule, but it is conceivable that different rules govern growth at different points in the trajectory. It may be possible to accurately approximate the rules governing the remodeling of networks over time, modelled either by changing heuristic estimates (e.g. changing of generative rules over time) or attempting to optimize a superordinate goal (e.g. computational efficiency and/or flexibility). Thirdly, our gene enrichment results are correlational, not causative. There remains an explanatory gap in determining whether and how these specific gene profiles support the sensitivity to connection formation. And crucially, the expression data are derived for a microarray analysis of post-mortem tissue samples from human adults^25^. The next steps will involve validating these findings in large scale developmental cohorts with available gene data, and forming casual links by applying GNMs to individuals with neurodevelopmental disorders of known genetic origin^46,47,52^.

In conclusion, we provide a unifying computational framework for conceptualizing the emergence of structural brain networks and their diversity. The emergence of brain networks can be understood as occurring via continual renegotiations of costs and values, but individuality emergences from their slightly different parameterization.

## METHODS

### Participants

Our sample were made up of children referred by practitioners working in specialist educational or clinical services to the Centre for Attention Learning and Memory (CALM), a research clinic at the MRC Cognition and Brain Sciences Unit, University of Cambridge (see Holmes, J. *et al*.^53^ for the full protocol of assessment, and refs^9–13^ for prior work using the same cohort). This cohort of children is intentionally heterogenous. Referrers were asked to identify children with cognitive problems related to learning, with primary referral reasons including difficulties with ongoing problems in “language”, “attention”, “memory”, or “learning / poor school progress”. Exclusion criteria were uncorrected problems in vision or hearing, English as a second language, or a causative genetic diagnosis. Children could have single, multiple or no formally diagnosed learning difficulty or neurodevelopmental disorder. The CALM cohort contains N=967 total children (N=805 referred; N=162 unreferred). Of these, N=299 undertook MRI scanning of which N=279 had usable MRI data (see *MRI acquisition and pre-processing*). N=270 of these had cognitive data available (see *Cognitive and learning assessments*). This sample includes 65.9% boys, mean age 117.8 months, age range was 66-223 months and 78 that came from the non-referred comparison sample. Demographic information of the N=270 subjects included in our sample is provided in **Supplementary Table 5**.

### MRI acquisition and pre-processing

Magnetic resonance imaging data were acquired at the MRC Cognition and Brain Sciences Unit in Cambridge, on the Siemens 3 T Tim Trio system (Siemens Healthcare) using a 32-channel quadrature head coil. N=299 CALM children underwent MRI scanning. 20 scans were not useable due to excessive motion (>3 mm movement during the diffusion sequence estimated through FSL eddy), leaving an MRI sample of N=279 children. T1-weighted volume scans were acquired using a whole brain coverage 3D Magnetization Prepared Rapid Acquisition Gradient Echo (MP RAGE) sequence acquired using 1 mm isometric image resolution. Echo time was 2.98ms, and repetition time was 2,250ms. Diffusion scans were acquired using echo-planar diffusion-weighted images with an isotropic set of 60 noncollinear directions, using a weighting factor of *b* = 1,000s × mm^-2^, interleaved with a T2-weighted (*b* = 0) volume. Whole brain coverage was obtained with 60 contiguous axial slices and isometric image resolution of 2 mm. Echo time was 90ms and repetition time was 8,400ms.

### Connectome construction

MRI scans were converted from the native DICOM to compressed NIfTI-1 format. Next, correction for motion, eddy currents, and field inhomogeneities was applied using FSL eddy. Furthermore, we submitted the images to nonlocal means de-noising^54^ using DiPy v0.11^55^ to boost signal-to-noise ratio. A constant single angle (CSA) model was fitted to the 60-gradient-direction diffusion-weighted images using a maximum harmonic order of 8 using DiPy. Whole-brain probabilistic tractography was performed with 8 seeds on all voxels. The step size was set to 0.5 and the maximum number of crossing fibers per voxel to 2. For ROI definition, T1-weighted images were submitted to nonlocal means denoising in DiPy, robust brain extraction using ANTs v1.9^56^, and reconstruction in FreeSurfer v5.3 (http://surfer.nmr.mgh.harvard.edu).

Regions of interest (ROIs) were based on the Desikan-Killiany parcellation of the MNI template^57^ with 34 cortical ROIs per hemisphere. To construct the connectivity matrix, the number of streamlines intersecting both ROIs was estimated and transformed into a density map for each pairwise combination of ROIs. A symmetric intersection was used so that streamlines starting and ending in each ROI were averaged. Self-connections were removed. To produce binarized connectomes from the resulting 68-by-68 streamline matrix, we enforced an average connectome density of ρ=10% (as in Betzel *et al*.^25^), resulting in a streamline threshold of 18 streamlines (i.e. a minimum of 18 streamlines must have connected two regions for us to consider the presence of an anatomical connection).

### Generative network models

Starting with a sparse seed network (25 bi-directional edges that were common across all *N* = 270 subjects), edges were added one at a time over a series of steps until a total number of connections were placed that equaled that of the target observed connectome (group level connections, mean = 231.4 and SD = 19.1). Each step allows for the possibility that any pair of unconnected nodes will be connected. Connections are formed probabilistically, where the relative probability of connection formation, between nodes *n^u^* and *n^v^*, is given by equation (1). We used thirteen previously studied non-geometric rules^24,25^ to produce energy landscapes. Topological parameters were computed using our own internally developed functions that originated from the Brain Connectivity Toolbox (https://sites.google.com/site/bctnet/)^58^.

To evaluate the fitness of synthetic networks and optimize models, we defined an energy function that measures how dissimilar a synthetic network is to the observed network as defined by Betzel, R.F. *et al*.^25^. This is given in equation (2). Initially, we ran simulations across a defined a parameter space of 10,000 evenly-spaced combinations of *η* ± 7 and *γ* ± 7, for each generative rule (**Fig. 2a-d**). This was to capture their respective energy landscapes and to estimate their relative effectiveness at generating plausible networks. We then computed a further a set of 50,000 simulations within a much narrower low-energy window (−3.606 ≤ *η* ≤ 0.354 and 0.212 ≤ *γ* ≤ 0.495) of the matching algorithm (**Fig. 2f**) for all subsequent analysis. This is because the matching algorithm attained the lowest-energy networks and therefore best approximated individual-level connectomes.

### Cognitive and learning assessments

A large battery of cognitive, learning, and behavioral measures was administered in the CALM clinic.^53^ Nine cognitive tasks were used in the current study. N=9 children did not have available cognitive data (of the N=279 MRI sample) and were therefore excluded, leaving the final sample N=270 children. These children had no missing data. For full details of the processing of cognitive data, see Siugzdaite, R. *et al*^13^.

The following measures of fluid and crystallized reasoning were included: Matrix Reasoning, a measure of fluid intelligence^59^ (Wechsler Abbreviated Scale of Intelligence, WASI); Peabody Picture Vocabulary Test^60^ (PPVT). Phonological processing was assessed using the Alliteration subtest of the Phonological Awareness Battery^61^ (PhAB). Verbal and visuo-spatial short-term and working memory were measured using Digit Recall, Dot Matrix, Backward Digit Recall, and Mr X subtests from the Automated Working Memory Assessment^62,63^ (AWMA). Learning measures (literacy and numeracy) were taken from the Wechsler Individual Achievement Test II^64^ (WIAT II) and the Wechsler Objective Numerical Dimensions^65^ (WOND), apart from 78 of controls for which we used multiple subtests from the Woodcock Johnson for Verbal ability^66^.

### Gene expression data

Regional microarray expression data were obtained from six post-mortem brains provided by the Allen Human Brain Atlas (http://human.brain-map.org/).^31,32^ These datasets were based on microarray analysis of post-mortem tissue samples from six human donors aged between 18 and 68 years with no known history of neuropsychiatric or neurological conditions.^26^ Data were imported from Arnatkevičiūtė, A. *et al*.^32^ Since only two of the six brains included samples from the right hemisphere, analyses were conducted on the left hemisphere only. Probes where expression measures do not exceed the background in >50% samples were removed and genes that did not have a corresponding RNA-seq measure were removed. Probes with a Spearman’s correlation <0.2 with RNA-seq data were removed, and a representative probe with the highest correlation to RNA-seq data was selected for each gene. Sample assignment was computed by applying a 2 mm distance threshold. In total, a mean of 37.8 ± 22.5 (SD) samples were assigned to each ROI (min= 5; max = 92)^32^.

The fully pre-processed gene data comprised of a 34 by 10,027 matrix (**Supplementary Fig. 7**) of microarray array gene expression data. These data were used for a subsequent PLS analysis (see *Statistics; PLS analysis*) and pathway enrichment analysis (see *Gene enrichment analysis and visualization*).

### Statistics

#### Predictions of spatial embedding

To assess the performance of the optimal matching GNMs to produce networks with spatial embedding of topological characteristics, we averaged across each subject’s best performing simulation (which achieved the lowest energy; descriptive statistics shown in the top row of **Supplementary Table 2**) to produce a single 68 (ROIs) by 1 vector for each measure. We did the same for their observed connectomes. **Figure 3** shows their linear correlations. In **Supplementary Figure 2a** we run the same process for local efficiency (not included in the energy equation). For **Supplementary Figure 2b-e** we correlate the same networks, but not for spatial embedding as these are global network measures outside of the energy equation. All network measures were calculated using functions from the Brain Connectivity Toolbox^58^.

#### Global associations of parameters with graphical and morphological measures

In **Figure 4** we perform a group level correlational analysis between *η* and *χ* and observed global graph theory and cortical morphology measures. In each case, the observed measures were averaged across the whole cortex.

#### Variability in the decomposed wiring equation

To determine where variability arises in the growth of the networks, we decomposed the wiring equation for each subject. This was achieved by first running the optimal wiring equation for each subject and taking their cost (a static Euclidean distance matrix), matching and wiring probability matrices at each step in the network growth model. For each subject, we took all edges that existed within the simulation and computed their mean and standard deviation (**Fig. 5c-e**) and then determined their coefficient of variation (**Fig. 5f** shows their distributions). To then explore within-connectome variability, we performed the same analysis but collapsing across subjects to determine how nodes (summed rows of the matrix) and edges (elements of the matrix) vary (**Fig. 5g**). From this section onward, subject 162 was removed from the analysis as they were the only subject to have an optimally performing *χ* that was positive, which biased results due to being an outlier (as parametrized values are calculated as *K^γ^*), leaving a sample of N=269.

#### PLS analysis

We used PLS regression to address two distinct aspects of the study. First, we used PLS to determine the latent components of the wiring equation and connectome features which best explain cognitive task performance (**Fig. 6a**). The p_corr_ significance value of each component was determined by permuting the cognitive data N=1000 times and comparing the observed covariance explained to the null distribution. **Figure 6b-c** show the correlation of predictor and response scores and response and predictor loadings of the significant PLS1 component (p_corr_ = 0.009 and p_corr_ = 0.049; in terms of covariance explained by the PLS1 latent components) for each analysis respectively. Second, we used PLS to identify the linear combinations of genes that best predicted average nodal costs and values each subject’s optimal simulation (as outlined previously). For each of the N=269 subjects, two PLS analyses were performed, providing 538 separate PLS analyses. To assess the significance of each gene in terms of its loading, we ran N=1000 permutations of the response variable for each PLS. This allowed us to compute a gene loading p_corr_ for each component of the PLS which was collapsed across subjects (as visualized in **Supplementary Fig. 3b**) for gene enrichment analysis (see *Gene enrichment analysis and visualization*).

### Gene enrichment analysis and visualization

We next aimed to elucidate the biological processes (BPs) and cellular components (CCs) for which our gene lists converged on. A BP is defined as representing a specific objective that the organism is genetically programmed to achieve. A BP is accomplished by a particular set of molecular functions carried out by specific gene products (or macromolecular complexes), often in a highly regulated manner and in a particular temporal sequence (https://www.ebi.ac.uk/QuickGO/term/GO:0008150) On the other hand, a CC is defined as a location, relative to cellular compartments and structures, occupied by a macromolecular machine when it carries out a molecular function. There are two ways in which the gene ontology describes locations of gene products: (1) relative to cellular structures (e.g., cytoplasmic side of plasma membrane) or compartments (e.g., mitochondrion), and (2) the stable macromolecular complexes of which they are parts (e.g., the ribosome) (https://www.ebi.ac.uk/QuickGO/term/GO:0005575).

To elucidate BPs and CCs across the sample, genes with a p_corr_<0.05 following permutation testing on each component were deemed significant. This provided an individual-level vector of genes that were significant for an individual for each of the nodal costs and nodal values PLS1. To collapse across subjects, genes were then ordered according to their frequency in being significantly associated with connectome growth across subjects for that component. The list stopped when genes were significant for less than 10% of the sample. For each subject, PLS1 provided an average of 581.5 significant genes (SD 101.4) for nodal costs and 437.6 significant genes (SD 167.4) for nodal values (**Supplementary Fig. 3a**). When collapsed across subjects as described, the nodal costs PLS1 had 1427 genes and the nodal values PLS1 had 1584 genes ordered in terms of importance, which were then submitted to a pathway enrichment analysis.

For all information as to the enrichment and visualization pipeline, please refer to Reimand, J. *et al*.^28^. In short, biological process gene ontology (GO) annotations are the most commonly used resource for pathway enrichment analysis. g:Profiler^67^ (https://biit.cs.ut.ee/gprofiler/gost) searches a collection of gene sets representing GO terms and, in the ordered test, repeats a modified Fisher’s exact test on incrementally larger sub-lists of the input genes and reports the sub-list with the strongest enrichment. Multiple-test correction is applied to produce an adjusted P value (p_adj_)^33,67^ (as visualized in **Supplementary Fig. 3c-d**, which can be accessed via the links presented in **Supplementary Table 4**). To visualize biological pathways, we used ‘EnrichmentMap’ within Cytoscape v3.8.0 (http://www.cytoscape.org)^33,68^. All default parameters were used. Pathways are shown as nodes (representing enriched biological processes) that are connected by edges if the pathways share genes. Nodes are colored by their p_adj_ and edges are sized on the basis of the number of genes shared by the connected pathways. To then identify clusters of themes, AutoAnnotate v1.3.3 was used before manually curating the suggested theme names to accurately reflect all pathways within each theme.

### Data availability

The datasets supporting the current study have not been deposited in a public repository because of restrictions imposed by NHS ethical approval, but are available from the corresponding author on request.

## ACKNOWLEDGEMENTS

The Centre for Attention Learning and Memory (CALM) research clinic at the MRC Cognition and Brain Sciences Unit in Cambridge (CBU) is supported by funding from the Medical Research Council of Great Britain to Duncan Astle., Susan Gathercole, Rogier Kievit, and Tom Manly. The clinic is led by Joni Holmes. Data collection is assisted by a team of PhD students and researchers at the CBU that includes Joe Bathelt, Sally Butterfield, Giacomo Bignardi, Sarah Bishop, Erica Bottacin, Lara Bridge, Annie Bryant, Elizabeth Byrne, Gemma Crickmore, Fánchea Daly, Edwin Dalmaijer, Tina Emery, Laura Forde, Delia Fuhrmann, Andrew Gadie, Sara Gharooni, Jacalyn Guy, Erin Hawkins, Agniezska Jaroslawska, Amy Johnson, Jonathan Jones, Silvana Mareva, Elise Ng-Cordell, Sinead O’Brien, Cliodhna O’Leary, Joseph Rennie, Ivan Simpson-Kent, Roma Siugzdaite, Tess Smith, Stepheni Uh, Francesca Woolgar, Mengya Zhang, Grace Franckel, Diandra Brkic, Danyal Akarca, Marc Bennet, and Sara Joeghan.

The authors wish to thank the many professionals working in children’s services in the southeast and east of England for their support and to the children and their families for giving up their time to visit the clinic.

We are particularly grateful for Joe Bathelt for his help in guiding us through the construction of the streamline connectomes.

Danyal Akarca is supported by the Medical Research Council Doctoral Training Programme and Cambridge Trust Vice Chancellor’s Award Scholarship. Duncan Astle is supported by Medical Research Council program grant MC-A0606-5PQ41 respectively. Edward Bullmore is an NIHR Senior Investigator. The opinions expressed in this publication are those of the authors and do not necessarily reflect the views of the funding agencies.

## AUTHOR CONTRIBUTIONS

All authors conceived the model and wrote the manuscript. All simulations, analysis and connectome construction were carried out by Danyal Akarca.

## COMPETING FINANCIAL INTERESTS

Edward Bullmore is a member of the Scientific Advisory Board for Sosei Heptares. Other authors declare no competing financial interests.

