## Supplementary Figures for "A generative network model of neurodevelopment"

| Rule | | | Energy | | $\eta$ | | $\gamma$ | |
|---|---|---|---|---|---|---|---|---|
| **Name** | **Class** | ***K(n^i,n^j)*** | **Mean** | **SD** | **Mean** | **SD** | **Mean** | **SD** |
| **Spatial** | Spatial | *1* | 0.337 | 0.020 | -3.940 | 0.397 | N/A | N/A |
| **Neighbours** | Homophily | $\sum_w a_{iw} a_{wj}$ | 0.165 | 0.020 | -1.995 | 0.483 | 0.338 | 0.045 |
| **Matching** | Homophily | $\frac{\|\Gamma i \backslash j \cap \Gamma j \backslash i\|}{\|\Gamma i \backslash j \cup \Gamma j \backslash i\|}$ | 0.141 | 0.021 | -1.647 | 0.490 | 0.357 | 0.022 |
| **C-Average** | Cluster | $\frac{Ci}{2} + \frac{Cj}{2}$ | 0.181 | 0.016 | -3.070 | 0.378 | -6.203 | 0.735 |
| **C-Minimum** | Cluster | $min[C_i, C_j]$ | 0.252 | 0.020 | -3.626 | 0.396 | -6.131 | 0.795 |
| **C-Maximum** | Cluster | $max[C_i, C_j]$ | 0.182 | 0.014 | -3.095 | 0.387 | -6.153 | 0.735 |
| **C-Difference** | Cluster | $\|C_i - C_j\|$ | 0.322 | 0.025 | -6.193 | 0.744 | -3.480 | 2.012 |
| **C-Product** | Cluster | $C_i C_j$ | 0.223 | 0.024 | -3.401 | 0.393 | -5.964 | 0.926 |
| **D-Average** | Degree | $\frac{ki}{2} + \frac{kj}{2}$ | 0.221 | 0.019 | -3.516 | 0.502 | 2.419 | 0.332 |
| **D-Minimum** | Degree | $min[k_i, k_j]$ | 0.279 | 0.016 | -4.749 | 0.749 | 0.474 | 0.010 |
| **D-Maximum** | Degree | $max[k_i, k_j]$ | 0.216 | 0.019 | -3.557 | 0.602 | 2.255 | 0.332 |
| **D-Difference** | Degree | $\|k_i - k_j\|$ | 0.229 | 0.018 | -3.836 | 0.669 | 1.689 | 0.577 |
| **D-Product** | Degree | $k_i k_j$ | 0.277 | 0.018 | -4.605 | 0.738 | 0.399 | 0.082 |

**Supplementary Table 1** Generative rules, their formulae and descriptive statistics across the broad parameter space ($\eta \pm 7$, $\gamma \pm 7$). Descriptive statistics of the $\eta$ and $\gamma$ required to achieve the best performing network for each subject, across the generative rules.

| Matching: Narrow energy window | | | | | | |
|---|---|---|---|---|---|---|
| Averaged over top N networks | Energy | | $\eta$ | | $\gamma$ | |
| | Mean | SD | Mean | SD | Mean | SD |
| N = 1 | 0.090 | 0.010 | -1.565 | 0.327 | 0.349 | 0.033 |
| N = 10 | 0.101 | 0.011 | -1.579 | 0.334 | 0.352 | 0.036 |
| N = 100 | 0.116 | 0.013 | -1.614 | 0.372 | 0.355 | 0.038 |
| N = 500 | 0.1326 | 0.015 | -1.658 | 0.427 | 0.357 | 0.040 |

**Supplementary Table 2** Descriptive statistics of high performing wiring parameters $\eta$ and $\gamma$ and the energy of the networks they produce. Parameters were selected using the matching generative rule with 50,000 simulations, evenly spaced across the narrow window $-3.606 < \eta < 0.354$ and $0.212 < \gamma < 0.495$. Parameters were averaged across a variable N number of high performing wiring combinations.

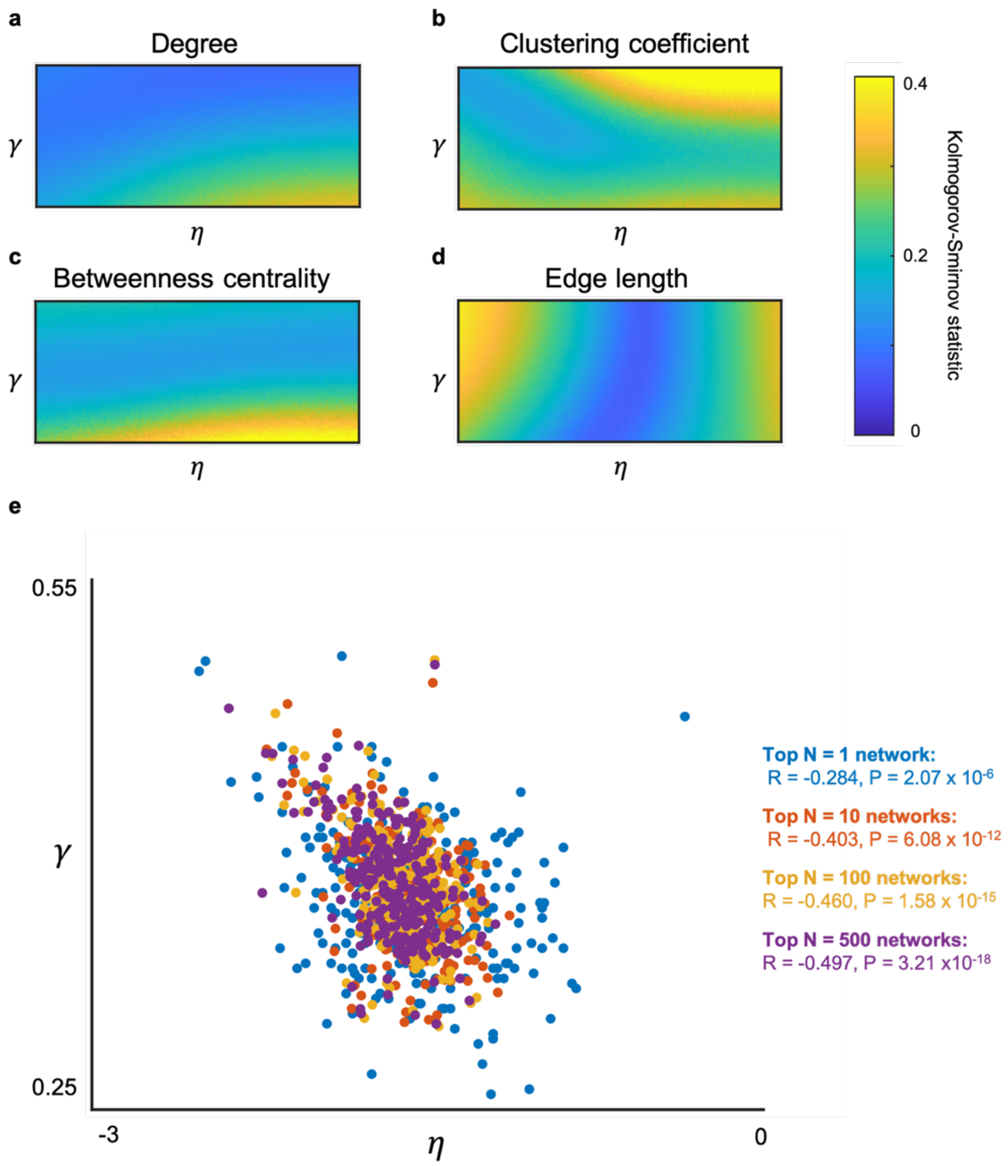


**Supplementary Figure 1** Spatial variation in KS statistics across the narrow matching window, and the trade-off between $\eta$ and $\gamma$ in the same space for variable N number of high performing wiring combinations. (**a**) Degree KS statistics (**b**) Clustering coefficient KS statistics (**c**) Betweenness-centrality KS statistics (**d**) Edge length statistics. (**e**) Within the narrow window, $\eta$ and $\gamma$ trade-off such that subjects with a larger $\gamma$ have a greater preponderance to have a larger magnitude negative $\eta$.

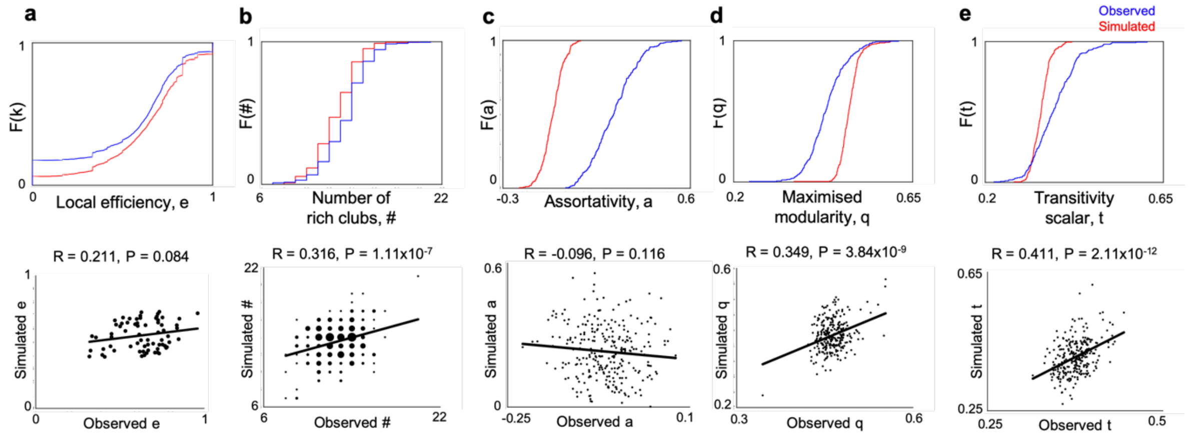


**Supplementary Figure 2** Cumulative density functions of topological and network features that were not included in the energy equation, and their predictions. Simulated measures were calculated from the top performing networks. (**a**) Local efficiency was calculated as in **Figure 3**, by averaging across subjects. Spatial embedding prediction is shown. Global measures for each subject are shown for the (**b**) Number of rich clubs. (**c**) Assortativity. (**d**) Modularity. (**e**) Transitivity.

| Global measures | $\eta$ | | | | | | | | $\gamma$ | | | | | | | |
|---|---|---|---|---|---|---|---|---|---|---|---|---|---|---|---|---|
| | Top N=1 simulated network | | Top N=10 simulated networks | | Top N=100 simulated networks | | Top N=500 simulated networks | | Top N=1 simulated network | | Top N=10 simulated networks | | Top N=100 simulated networks | | Top N=500 simulated networks | |
| | R | P | R | P | R | P | R | P | R | P | R | P | R | P | R | P |
| **Degree** | 0.0623 | 0.3074 | -0.0828 | 0.1748 | **-0.1209** | **0.0472** | **-0.1845** | **0.0023** | **0.3215** | **$6.5867x10^{-8}$** | **0.5765** | **$2.5715x10^{-25}$** | **0.5860** | **$2.7400x10^{-26}$** | **0.5947** | **$3.2074x10^{-27}$** |
| **Clustering coefficient** | **-0.2424** | **$5.7241x10^{-5}$** | **-0.4312** | **$1.1915x10^{-13}$** | **-0.5754** | **$3.3105x10^{-25}$** | **-0.6338** | **$9.7869x10^{-32}$** | **0.5879** | **$1.7144x10^{-26}$** | **0.7878** | **$2.3807x10^{-58}$** | **0.8314** | **$2.4001x10^{-70}$** | **0.8285** | **$1.8709x10^{-69}$** |
| **Betweenness centrality** | **-0.1595** | **0.0087** | -0.1128 | 0.0641 | **-0.1741** | **0.0041** | **-0.1391** | **0.0223** | **-0.2194** | **$2.8060x10^{-4}$** | **-0.4125** | **$1.6248x10^{-12}$** | **-0.4225** | **$4.0817x10^{-13}$** | **-0.4424** | **$2.2968x10^{-14}$** |
| **Edge length** | **0.3415** | **$8.4813x10^{-9}$** | **0.4752** | **$1.2926x10^{-16}$** | **0.5477** | **$1.5660x10^{-22}$** | **0.4925** | **$6.6608x10^{-18}$** | 0.1168 | 0.0553 | **0.2094** | **$5.3257x10^{-14}$** | **0.2287** | **$1.5018x10^{-4}$** | **0.2491** | **$3.4712x10^{-5}$** |
| **Efficiency** | 0.1049 | 0.0854 | 0.041 | 0.5020 | 0.0154 | 0.8005 | -0.0460 | 0.4516 | **0.2070** | **$6.2060x10^{-4}$** | **0.3988** | **$9.9888x10^{-12}$** | **0.4158** | **$1.0312x10^{-12}$** | **0.4340** | **$7.8898x10^{-14}$** |
| **Number of rich clubs** | 0.0861 | 0.1581 | 0.0177 | 0.7718 | 0.0090 | 0.8827 | 0.0206 | 0.7360 | **0.2041** | **$7.4004x10^{-4}$** | **0.3560** | **$1.7377x10^{-9}$** | **0.3832** | **$7.1277x10^{-11}$** | **0.4001** | **$8.4373x10^{-12}$** |
| **Assortativity** | 0.0130 | 0.8313 | 0.0358 | 0.5578 | 0.0311 | 0.6104 | 0.0531 | 0.3845 | **-0.1850** | **0.0023** | **-0.1964** | **0.0012** | **-0.1564** | **0.0101** | **-0.1602** | **0.0083** |
| **Transitivity scalar** | **-0.2548** | **$2.2667x10^{-5}$** | **-0.3989** | **$9,7813x10^{-12}$** | **-0.5281** | **$8.5927x10^{-21}$** | **-0.5767** | **$2.4941x10^{-25}$** | **0.4863** | **$1.9646x10^{-17}$** | **0.6383** | **$2.6553x10^{-32}$** | **0.6792** | **$7.0291x10^{-38}$** | **0.6689** | **$2.1791x10^{-36}$** |
| **Maximised modularity** | **-0.2249** | **$1.9449x10^{-4}$** | **-0.1747** | **0.0040** | **-0.2626** | **$1.234x10^{-5}$** | **-0.2374** | **$8.1647x10^{-5}$** | -0.1052 | 0.0844 | **-0.2291** | **$1.4583x10^{-4}$** | **-0.2432** | **$5.3864x10^{-5}$** | **-0.2543** | **$2.3398x10^{-5}$** |
| **Grey volume** | -0.0040 | 0.9482 | -0.1048 | 0.0856 | **-0.1644** | **0.0068** | **-0.1799** | **0.0030** | **0.1492** | **0.0141** | **0.2363** | **$8.8556x10^{-5}$** | **0.2278** | **$1.5917x10^{-4}$** | **0.2217** | **$2.4095x10^{-4}$** |
| **Surface area** | 0.0061 | 0.9207 | -0.0950 | 0.1196 | **-0.1461** | **0.0163** | **-0.1627** | **0.0074** | **0.1329** | **0.0291** | **0.2214** | **$2.4517x10^{-4}$** | **0.2141** | **$3.9639x10^{-4}$** | **0.2081** | **$5.7950x10^{-4}$** |
| **Cortical thickness** | -0.0331 | 0.5885 | -0.0909 | 0.1362 | **-0.1730** | **0.0044** | **-0.1979** | **0.0011** | **0.1262** | **0.0382** | **0.2058** | **$6.6834x10^{-4}$** | **0.1938** | **0.0014** | **0.1871** | **0.0020** |

**Supplementary Table 3** Correlational statistics between $\eta$ and $\gamma$ and observed global network and graphical measures. Global measures were calculated by averaging across the cortex. Parameters were averaged across a variable N number of best performing networks to determine how this influences associations. The first four rows (blue) correspond to measures minimised within the energy equation (2). The next five rows (green) correspond to measures not included in the energy equation. The final three rows (yellow) are cortical morphology measures which were not included in the energy equation. For all N, both $\eta$ and $\gamma$ are associated with a range of observed characteristics within and outside of the energy equation. Boldened numbers are associations which are significant at P<0.05.

**a**

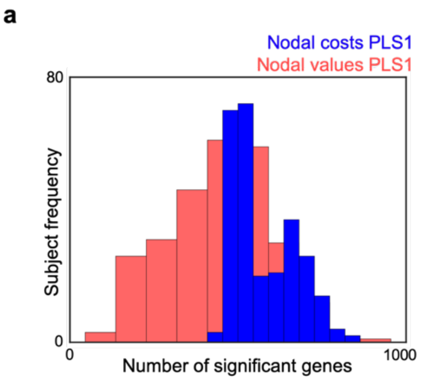


**b**

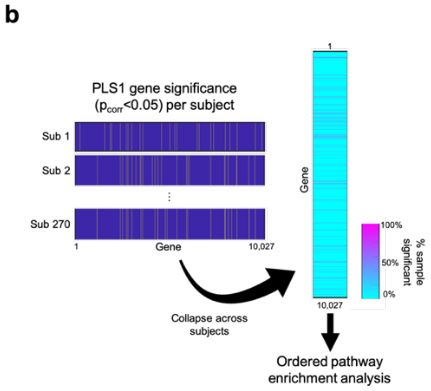


**c**

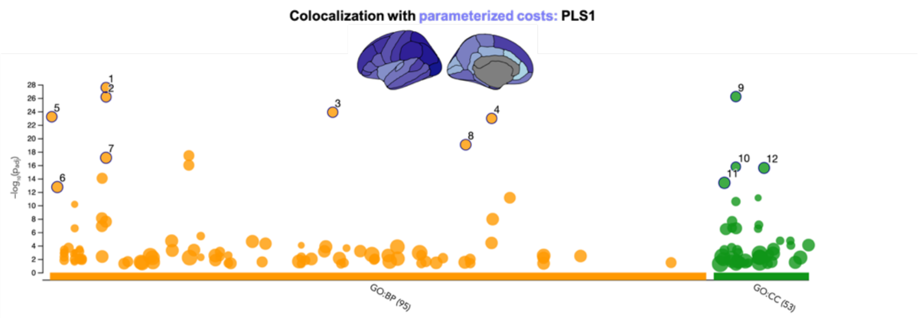


| ID | Source | Term ID | Term Name | $p_{adj}$ (query_1) |
|---|---|---|---|---|
| 1 | GO:BP | GO:0006614 | SRP-dependent cotranslational protein targeti... | $2.672 \times 10^{-28}$ |
| 2 | GO:BP | GO:0006613 | cotranslational protein targeting to membrane | $6.844 \times 10^{-27}$ |
| 3 | GO:BP | GO:0045047 | protein targeting to ER | $1.281 \times 10^{-24}$ |
| 4 | GO:BP | GO:0072599 | establishment of protein localization to endopl... | $1.060 \times 10^{-23}$ |
| 5 | GO:BP | GO:0000184 | nuclear-transcribed mRNA catabolic process, ... | $6.060 \times 10^{-24}$ |
| 6 | GO:BP | GO:0000956 | nuclear-transcribed mRNA catabolic process | $1.799 \times 10^{-13}$ |
| 7 | GO:BP | GO:0006612 | protein targeting to membrane | $7.750 \times 10^{-18}$ |
| 8 | GO:BP | GO:0070972 | protein localization to endoplasmic reticulum | $8.765 \times 10^{-20}$ |
| 9 | GO:CC | GO:0022626 | cytosolic ribosome | $6.098 \times 10^{-27}$ |
| 10 | GO:CC | GO:0022625 | cytosolic large ribosomal subunit | $1.689 \times 10^{-16}$ |
| 11 | GO:CC | GO:0005840 | ribosome | $4.218 \times 10^{-14}$ |
| 12 | GO:CC | GO:0044391 | ribosomal subunit | $2.533 \times 10^{-16}$ |

**d**

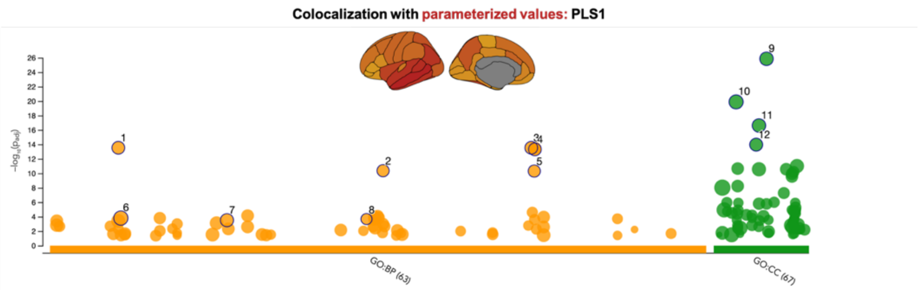


| ID | Source | Term ID | Term Name | $p_{adj}$ (query_1) |
|---|---|---|---|---|
| 1 | GO:BP | GO:0007268 | chemical synaptic transmission | $2.930 \times 10^{-14}$ |
| 2 | GO:BP | GO:0050804 | modulation of chemical synaptic transmission | $4.298 \times 10^{-11}$ |
| 3 | GO:BP | GO:0098916 | anterograde trans-synaptic signaling | $2.930 \times 10^{-14}$ |
| 4 | GO:BP | GO:0099536 | synaptic signaling | $4.787 \times 10^{-14}$ |
| 5 | GO:BP | GO:0099177 | regulation of trans-synaptic signaling | $4.859 \times 10^{-11}$ |
| 6 | GO:BP | GO:0007399 | nervous system development | $1.550 \times 10^{-4}$ |
| 7 | GO:BP | GO:0031175 | neuron projection development | $3.060 \times 10^{-4}$ |
| 8 | GO:BP | GO:0048167 | regulation of synaptic plasticity | $2.111 \times 10^{-4}$ |
| 9 | GO:CC | GO:0045202 | synapse | $1.389 \times 10^{-26}$ |
| 10 | GO:CC | GO:0030054 | cell junction | $1.281 \times 10^{-20}$ |
| 11 | GO:CC | GO:0043005 | neuron projection | $2.379 \times 10^{-17}$ |
| 12 | GO:CC | GO:0036477 | somatodendritic compartment | $1.080 \times 10^{-14}$ |

**Supplementary Figure 3** Gene loading analysis and pathway enrichment (**a**) Distributions of significant genes ($p_{corr} < 0.05$) for the nodal-costs and nodal-values PLS1, which provided an average of 581.5 significant genes and 437.6 significant respectively across the sample. (**b**) To collapse across subjects, we ranked genes according to the number of subjects for which each gene was significant and submitted genes above 10% to an ordered pathway enrichment analysis. (**c**) Pathway enrichment analysis summarises large gene sets as a smaller list of more easily interpretable pathways that can be visualised to identify main biological themes. Outcomes of Gene Ontology Biological Processes (BP) from the g:Profiler statistical enrichment analysis. We highlight a number of highly significant results.

| PLS | Response | Number of genes | Gene Ontology | Link |
|---|---|---|---|---|
| PLS1 | Parametrised costs | 1427 | Biological processes and cellular components | https://biit.cs.ut.ee/gplink/l/TYGoJHihSs |
| | Parameterised values | 1584 | Biological processes and cellular components | https://biit.cs.ut.ee/gplink/l/H_gK8UIVR1 |

**Supplementary Table 4** Gene queries submitted to g:Profiler. Ordered gene lists were computed by taking subject-specific significant ($p_{corr}<0.05$) genes according to their frequency across the sample, up to 10% of the sample.

| N | Age (months) | | Sex | | Referral status | |
|---|---|---|---|---|---|---|
| | Mean | SD | Male | Female | Referred | Non-referred |
| 270 | 117.8 | 25.8 | 178 | 92 | 192 | 78 |

**Supplementary Table 5** Demographic information of the sample.
